# Glutamine-addiction in cisplatin resistant cancer cells is mediated by SLC7A11 and can be targeted with asparaginase therapy

**DOI:** 10.1101/2024.07.19.604261

**Authors:** Jiantao Wang, Robert Strauss, Jiri Bartek, Sean G Rudd

## Abstract

Linking disease phenotypes with molecular targets is key to the rational design of treatment interventions. Resistance to the chemotherapeutic cisplatin is one of the major factors limiting the clinical utility of this therapy, which is central to the treatment of a variety of solid malignancies. In this study, we couple the upregulation of a chemoresistant factor, the glutamate-cystine antiporter SLC7A11, with the addiction of cisplatin-resistant cancer cells to extracellular glutamine. In doing so, we thus provide a putative biomarker for this acquired metabolic dependency of chemoresistance. Subsequently, we evaluate various therapeutic strategies to selectively kill SLC7A11^high^ cisplatin-resistant cancer cells, identifying cross-resistance to ferroptosis-inducing compounds and hypersensitivity to glutaminase inhibitor CB-839. We identify enzymatic depletion of extracellular glutamine using the long-standing anti-leukemic therapy asparaginase (ASNase), which possesses glutaminase activity, as a potential approach, and show this can be successfully combined with cisplatin in cell models. In summary, this study mechanistically links an acquired metabolic dependency of chemoresistant cancer cells with a putative biomarker and provides a potentially actionable strategy to target these drug resistant cells warranting further investigation.

## Introduction

Conventional chemotherapeutic agents remain a fundamental treatment modality for patients with cancer [1]. The platinum derivative cisplatin (cis-diamminedichloroplatinum; CDDP) is first line treatment for a variety of solid malignancies, including cancers of the lung, bladder, and ovaries, amongst many others. Whilst a good initial response to this agent is common, the duration of this response varies greatly and eventual disease relapse can occur, which is a major clinical problem [2]. Thus, gaining a molecular understanding of chemoresistance and using this knowledge to design actionable therapeutic strategies to overcome it is vital.

The molecular mode-of-action of cisplatin and platinum-derived compounds is principally understood to centre around crosslinking of DNA strands, which are cytotoxic DNA lesions that prevent essential metabolic processes such as replication and transcription [3,4]. To do this, cisplatin, a prodrug, is first activated inside cancer cells by aquation, which occurs spontaneously in the cytoplasm owing to lower intracellular chloride concentrations compared to the extracellular milieu. This produces a potent electrophile that can react with biological molecules in both the cytosol and nucleus, including the bases in the DNA duplex [3,4]. Following cytotoxic DNA damage, DNA damage signalling will promote induction of the mitochondrial apoptotic pathway resulting in cell death, which can also be supported by cytosolic targets of aquated-cisplatin [5].

Given this multi-step mode-of-action of cisplatin and platinum-derived agents, multiple resistance mechanisms exist, which have been widely documented in both the clinical and preclinical setting [2,3,5–7]. These include reduced drug uptake or increased drug efflux, elevated DNA repair proficiency to remove cytotoxic cisplatin adducts, or alternatively increased tolerance of cisplatin adducts and failure to induce apoptotic cell death. In addition, aquated-cisplatin can be de-toxified from cells by reacting with thiol containing molecules, such as glutathione (GSH).

GSH is a tripeptide consisting of *γ*-glutamate, cysteine and glycine, with a major role in maintaining redox homeostasis in cells by reacting with harmful by-products of aerobic metabolism [8]. Owing to the nucleophile thiol residue which lacks steric hindrance, and being highly abundant inside cells, the thiol in GSH can react with electrophiles such as aqauted-cisplatin, thereby protecting cells of this cytotoxic compound. The cysteine component of GSH is generated from cystine, which is imported by antiporter SLC7A11 at the expense of exporting glutamate. The glutamate component of GSH is derived from glutamine, a non-essential amino acid that is highly abundant in the human body [9]. Glutamine can be obtained through diet and can also be synthesised inside cells by glutamine synthetase (GS). Extracellular glutamine is imported into cells by soluble carrier transporters such as ASCT2 (SLC1A5). Within cells, glutamine is involved in many metabolic processes, including nucleotide biosynthesis, energy production via the TCA cycle, and generation of the antioxidant GSH to maintain redox homeostasis [9].

Several studies have reported that cancer models with acquired resistance to cisplatin become dependent upon glutamine [10–16], and a number of distinct mechanisms have been described. For instance, elevated expression of glutamine transporter ASCT2 and glutaminase (GLS) has been documented in platinum-resistant ovarian cancer cell models, reported to confer resistance via increased use of glutamine in the TCA cycle [10]. A similar mechanism was recently reported in a chondrosarcoma cell model [16]. Glutamine consumption via GLS was also demonstrated to contribute to acquired cisplatin resistance in HeLa cells [14]. Another study in a cisplatin-resistant ovarian cancer cell line identified methylation-mediated silencing of GS to increase GSH production as cause of resistance [11]. Alternatively, increased use of glutamine to fuel nucleotide biosynthesis has also been documented, rendering cisplatin-resistant models sensitive to antimetabolite therapies [12].

In the present study, we sought to further investigate the mechanisms of glutamine dependence using cell models with acquired resistance to cisplatin and evaluate therapeutic strategies to selectively target these cells. In doing so, we identified upregulation of SLC7A11 as an additional mechanism of glutamine dependence in cisplatin-resistant models, and subsequently demonstrate the applicability of enzymatically depleting extracellular glutamine using the approved therapy asparaginase (ASNase), as one such approach.

## Results

### Acquired resistance to cisplatin can lead to dependency upon extracellular glutamine

To begin our study, we first assembled a collection of cancer cell lines with acquired resistance to cisplatin, consisting of those derived from the lung adenocarcinoma cell line HCC4006, the bladder cancer cell line NTUB1, and the ovarian cancer cell line A2780. The parental cell lines and their cisplatin-resistant counterparts were subjected to a 3-day incubation of a cisplatin dose-response in a proliferation inhibition assay, which confirmed acquired resistance to this therapy (**Fig 1a**). Subsequently we assessed the glutamine dependence of these cell models by plating cells in complete medium in the presence and absence of glutamine and following 2 days measuring remaining viable cells, which demonstrated that the cisplatin resistant HCC4006 and NTUB1 cells, but not A2780, were more dependent upon extracellular glutamine for cell proliferation than their parental counterparts (**Fig 1b**). Furthermore, we could recapitulate this result in a colony outgrowth assay performed in the presence or absence of glutamine (**Fig 1c**).

**Figure 1.**
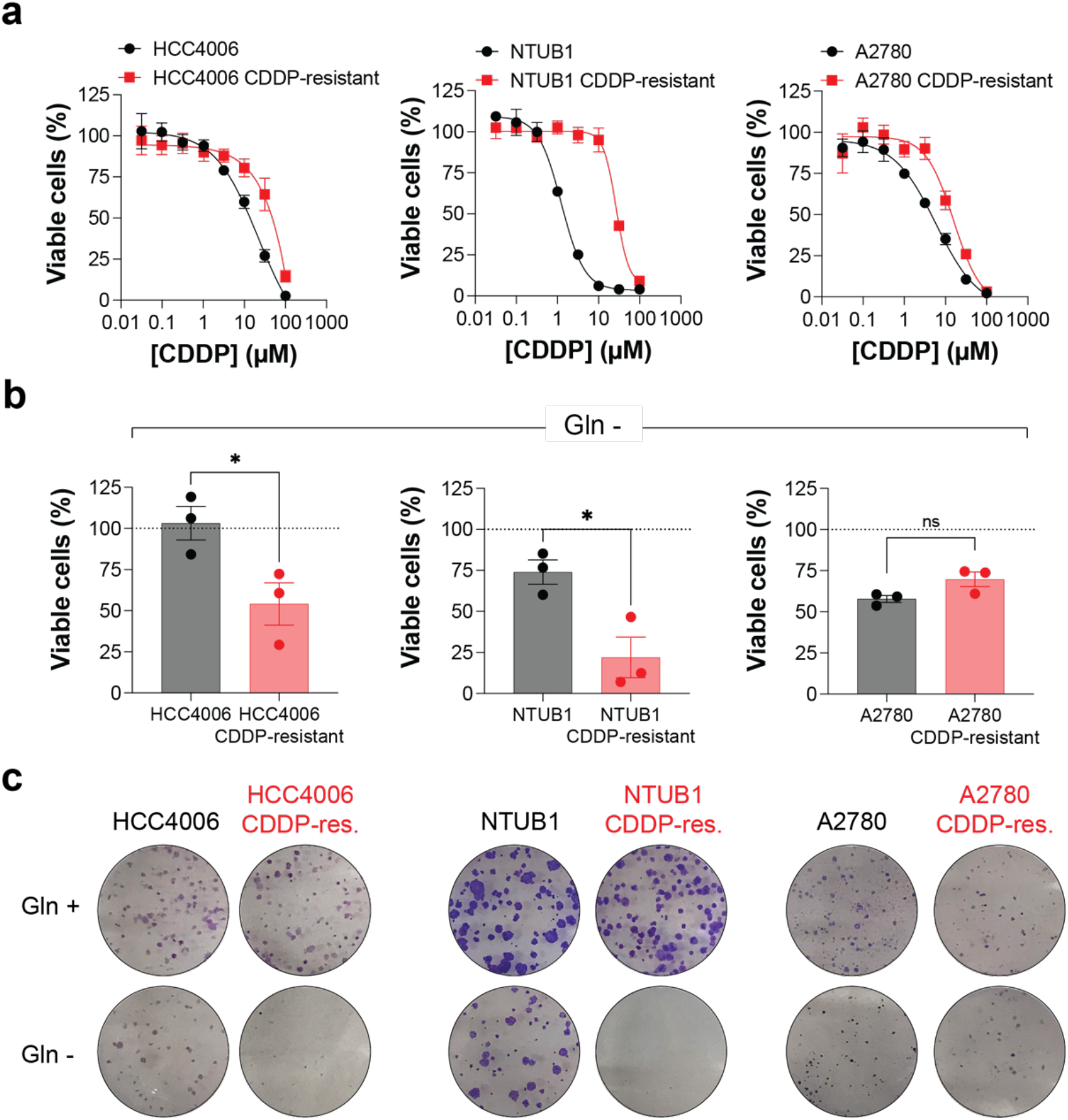
Acquired resistance to cisplatin can lead to a dependency upon extracellular glutamine in cell line models. **a)** Parental and cisplatin (CDDP)-resistant cell line pairs were exposed to a CDDP dose-response and cell viability measured after 3 days via resazurin reduction. Mean values of independent experiments (HCC4006 n=3; NTUB1 n=2, A2780 n=2) each performed in duplicate shown, error bars indicate SEM. **b)** Parental and CDDP-resistant cell line pairs were cultured in complete media or media lacking glutamine for 2 days prior to resazurin reduction assay. Proportion of viable cells in the absence of glutamine relative to cells cultured in complete media (dashed line) plotted. Mean values from independent experiments shown, each performed with a minimum of 6 technical replicates, bars and error bars indicate mean and SEM (n=3). Unpaired two-tailed t-test: *, P > 0.05; ns, not significant. **c)** Parental and CDDP-resistant cell line pairs were cultured in complete media or media lacking glutamine in a colony outgrowth assay before fixation and staining. Representative images shown (n=2).

### Elevated SLC7A11 contributes to glutamine dependence of cisplatin-resistant cancer cells

Seeking to understand the mechanistic basis of glutamine dependence, we next performed immunoblot analysis of lysates from the cell line pairs to determine the protein levels of enzymes involved in glutamine metabolism (**Fig. 2a**). Protein levels of glutamine importer ASCT2 and glutaminases GLS1/2 were unchanged between the parental and cisplatin-resistant cell line pairs, however glutamine synthetase (GS) and the glutamate-cystine antiporter SLC7A11 showed differential expression (**Fig 2b**). Specifically, cisplatin-resistant NTUB1 and A2780 cell lines had no detectable GS protein, whilst cisplatin-resistant HCC4006 and NTUB1 had elevated protein levels of SLC7A11 (**Fig 2b**). Given the glutamine-dependent cisplatin-resistant cell lines (HCC4006 and NTUB1) both up-regulate SLC7A11, we then interrogated whether the glutamine-dependence is a consequence of this up-regulation. In the cisplatin-resistant cell models we depleted SLC7A11 protein using siRNA, confirmed by immunoblot analysis (**Fig 2c**), and subjected these cells to glutamine-withdrawal before measuring remaining viable cells with orthogonal readouts. In both instances, we observed that depletion of SLC7A11 partially reduced the glutamine-dependence of the cisplatin-resistant cancer models (**Fig 2d, e**), supporting that up-regulation of SLC7A11 contributes to their glutamine dependency.

**Figure 2.**
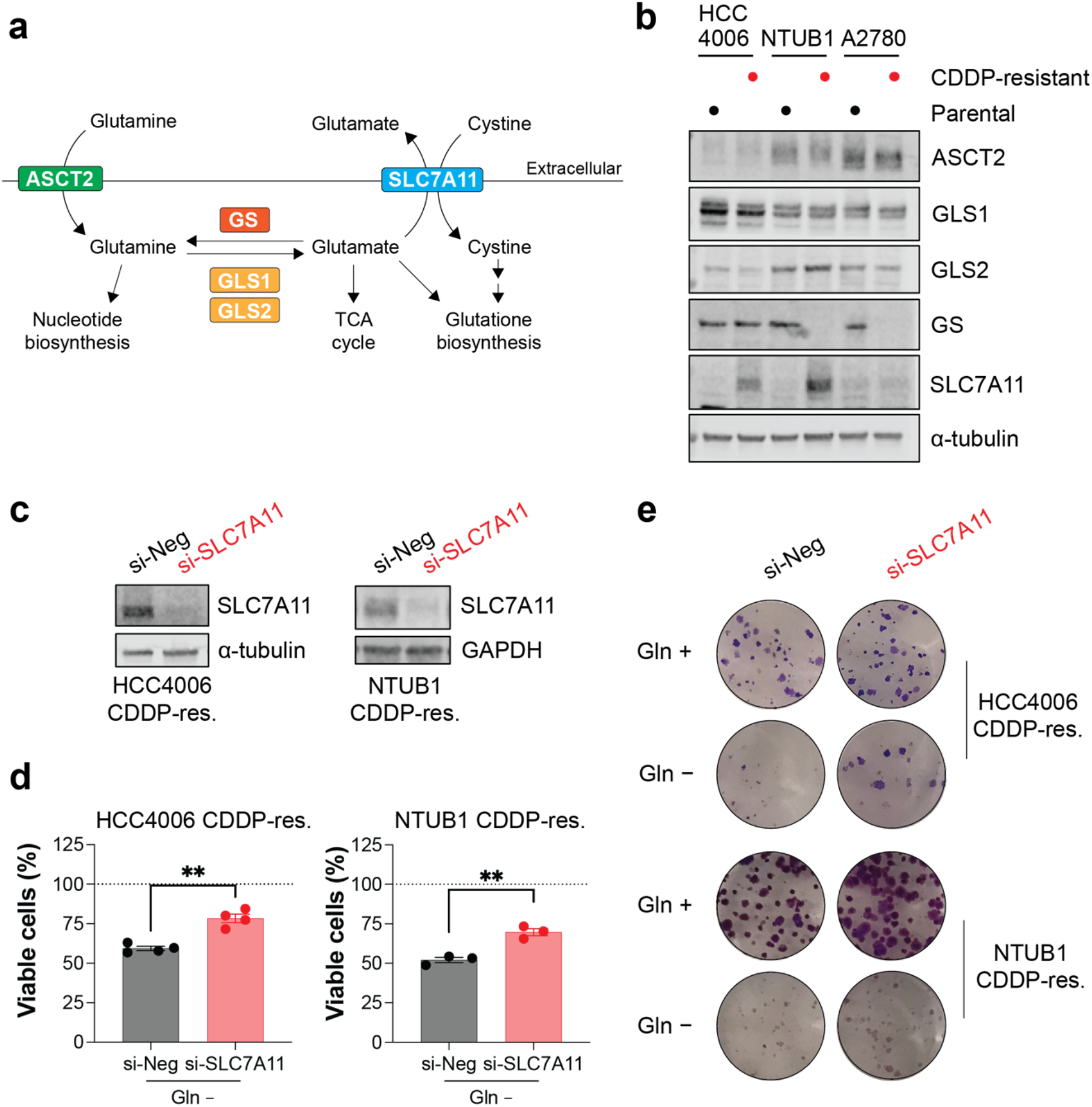
Elevated SLC7A11 contributes to glutamine dependence of cisplatin-resistant cancer cell models. **a)** Schematic of glutamine metabolism. ASCT2, glutamine importer; GLS1 and 2, glutaminase 1 and 2; GS, glutamine synthetase; SLC7A11, cystine glutamate antiporter. **b)** Immunoblot analysis of lysates prepared from parental and cisplatin (CDDP)-resistant pairs with the indicated antibodies. Representative images of cropped blots shown (n=2). **c)** HCC4006 or NTUB1 parental and CDDP-resistant cell line pairs were transfected with control (si-Neg) or SLC7A11 targeting (si-SLC7A11) siRNA and after 3 days, lysates prepared and analysed by immunoblot with the indicated antibodies. Representative images of cropped blots shown (n=2) **d)** Cells as transfected in (c) were cultured in complete media or media lacking glutamine for 2 days before resazurin reduction assay. Proportion of viable cells in the absence of glutamine relative to cells cultured in complete media (dashed line) plotted. Mean values from independent experiments shown, each performed with a minimum of 6 technical replicates, bars and error bars indicate mean and SEM (n=3-4). Unpaired twotailed t-test: **, P > 0.01. **e)** Cells as transfected in (c) were cultured in media lacking glutamine in a colony outgrowth assay before fixation and staining. Representative images shown (n=2).

### SLC7A11^high^ cisplatin-resistant cancer cells are cross-resistant to ferroptosis inducers and hypersensitive to glutaminase inhibition

Having identified a sub-population of cisplatin-resistant cancer cells, i.e., those with high expression of SLC7A11 (SLC7A11^high^) that are dependent upon extracellular glutamine, represented here by cisplatin-resistant HCC4006 and NTUB1, we next investigated the utility of various treatment interventions to target these cells. We focused upon treatments targeting glutamine metabolism or subsequent downstream processes.

We began by evaluating the cell line panel for sensitivity to erastin, targeting SLC7A11 [17], and RSL-3, identified as targeting the lipid peroxidase GPX4 [18] although this was recently called into question [19]. Regardless, these molecules induce ferroptosis, an iron-dependent cell death mechanism with broad implications for overcoming therapy resistance [20]. Immunoblot analysis of the cell line panel indicated that SLC7A11^high^ cisplatin-resistant cancer cell lines downregulate GPX4 (**Fig 3b**). In the context of acquired cisplatin-resistance, we observed that SLC7A11^high^ cisplatin-resistant cancer cells were differentially sensitive to erastin with HCC4006-cisplatin resistant cells becoming cross-resistant to this molecule, whilst no difference was observed in NTUB1-cisplatin resistant cells (**Fig 3c**). However, with regards to RSL-3, both SLC7A11^high^ cisplatin-resistant cell models became cross-resistant, whilst enhanced sensitivity was observed in the A2780-cisplatin resistant cell model (**Fig 3c**).

**Figure 3.**
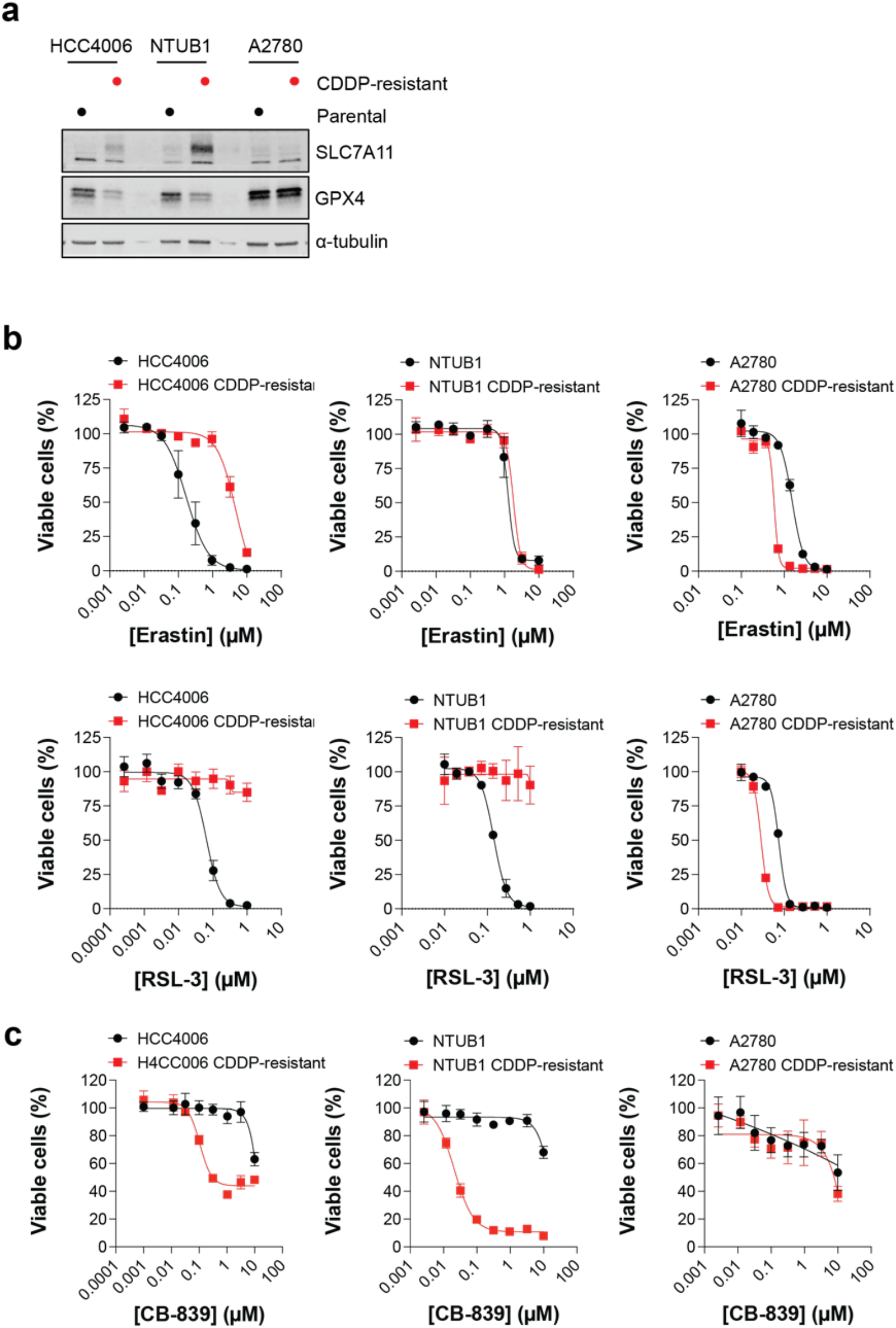
SLC7A11^high^ cisplatin-resistant cells are cross-resistant to ferroptosis inducers and hypersensitive to glutaminase inhibition. **a)** Immunoblot analysis of lysates prepared from parental and cisplatin (CDDP)-resistant pairs with the indicated antibodies. Representative images of cropped blots shown (n=2). **b, c**) Parental and CDDP-resistant cell line pairs were exposed to a dose-response of the indicated compound and cell viability measured after 3 days via resazurin reduction. Mean values of independent experiments (n=2-3) each performed in duplicate shown, error bars indicate SEM.

CB-839 (telaglenastat) is a potent and selective, orally bioavailable GLS1 inhibitor in advanced clinical testing [21,22] (**Fig 3a**). GLS1 and GLS2 showed little differential expression amongst the cisplatin resistant and parental cell line pairs (**Fig 2b**). Treatment of the cell line panel with a doseresponse of this compound revealed a striking sensitivity of SLC7A11^high^ cisplatin-resistant cancer cells, with the half maximal inhibitory concentration (IC_50_) values decreasing by several orders of magnitude compared to parental counterparts (**Fig 3d**). In contrast, A2780-cisplatin resistant cell model and parental counterpart showed no differential sensitivity (**Fig 3d**). Altogether, these data demonstrate that acquired resistance to cisplatin can result in differential sensitivity to several treatment interventions.

### SLC7A11^high^ cisplatin-resistant cancer cells are sensitive to ASNase therapy

Next, we evaluated the decades old anti-leukemic therapy ASNase. Although the clinical use of ASNase centres upon the enzymatic hydrolysis of extracellular asparagine, required for the proliferation of asparagine auxotroph leukemic cells, ASNase also possesses glutaminase activity, and accordingly also enzymatically depletes extracellular glutamine [23,24] (**Fig 4a**). Thus, here, we interrogated the applicability of this approach to target glutamine addicted SLC7A11^high^ cisplatin resistant cancer cells. Supplementation of cell medium with ASNase reduced the proportion of viable cells, with an increased sensitivity observed in the SLC7A11^high^ cisplatin resistant models (**Fig 4c**). In the colony formation assay, this specific sensitivity of the SLC7A11^high^ cisplatin resistant cells was much more pronounced (**Fig 4d**).

**Figure 4.**
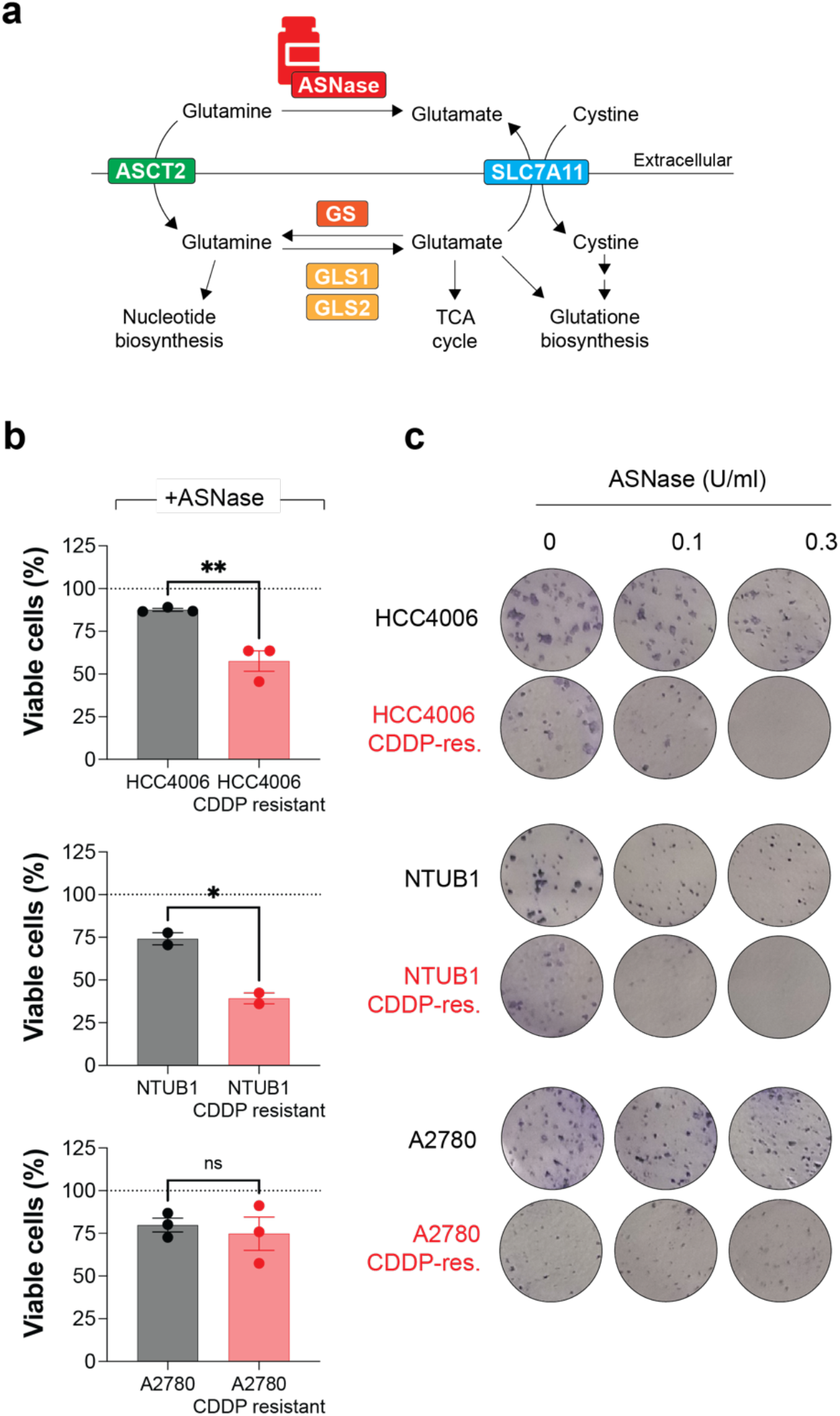
SLC7A11^high^ cisplatin-resistant cancer cells are sensitive to ASNase therapy. **a)** Schematic of glutamine metabolism indicating ASNase can target extracellular glutamine. **b)** Parental and cisplatin (CDDP)-resistant cell line pairs were cultured in complete media or media supplemented with ASNase (0.3 U/ml) for 3 days before resazurin reduction assay. Proportion of viable cells in the absence of ASNase relative to cells cultured in complete media (dashed line) plotted. Mean values from independent experiments shown, each performed in duplicate, bars and error bars indicate mean and SEM (n=2-3). Unpaired two-tailed t-test: **, P > 0.01; *, P > 0.05; ns, not significant. **c)** Parental and CDDP-resistant cell line pairs were cultured in complete media or media supplemented with the indicated doses of ASNase in a colony outgrowth assay before fixation and staining. Representative images shown (n=2).

### ASNase and cisplatin combination treatment in SLC7A11^high^ cisplatin-resistant cancer cells results in additive inhibition of proliferation

Given ASNase is a cancer therapy in routine clinical practice, we sought to evaluate the applicability of this approach further, and test whether it can be successfully combined with cisplatin therapy. We speculated that enzymatic depletion of extracellular glutamine by ASNase may re-sensitise the cisplatin-resistant cells to cisplatin. In the colony outgrowth experiments, evaluating each drug alone and in combination, we observed that addition of ASNase in combination with cisplatin in the SLC7A11^high^ cisplatin-resistant cell lines (HCC4006 and NTUB1) led to a further reduction in colony outgrowth (**Fig 5a**), suggestive of at least an additive effect in reducing cell proliferation. To assess this combination in more depth, SLC7A11^high^ cisplatin-resistant cancer cells were seeded upon dose-response matrices of cisplatin and ASNase and cell viability measured following a 3-day treatment. ASNase alone dose-dependently decreased the proportion of viable cells, and furthermore, sensitised these cell models to cisplatin in a dosedependent manner (**Fig 5b, c**). We next used the data generated to assess the drug combination effect over the entire drug matrix and obtained dose-response landscapes consistent with additive cell killing, although peaks of synergistic cell killing could be observed at select doses (**Fig 5d**). Analysis of this data by several drug combination models indicated the combination of cisplatin and ASNase in SLC7A11^high^ cisplatin-resistant cell lines was largely additive in reducing cancer cell proliferation (**Fig 5e**).

**Figure 5.**
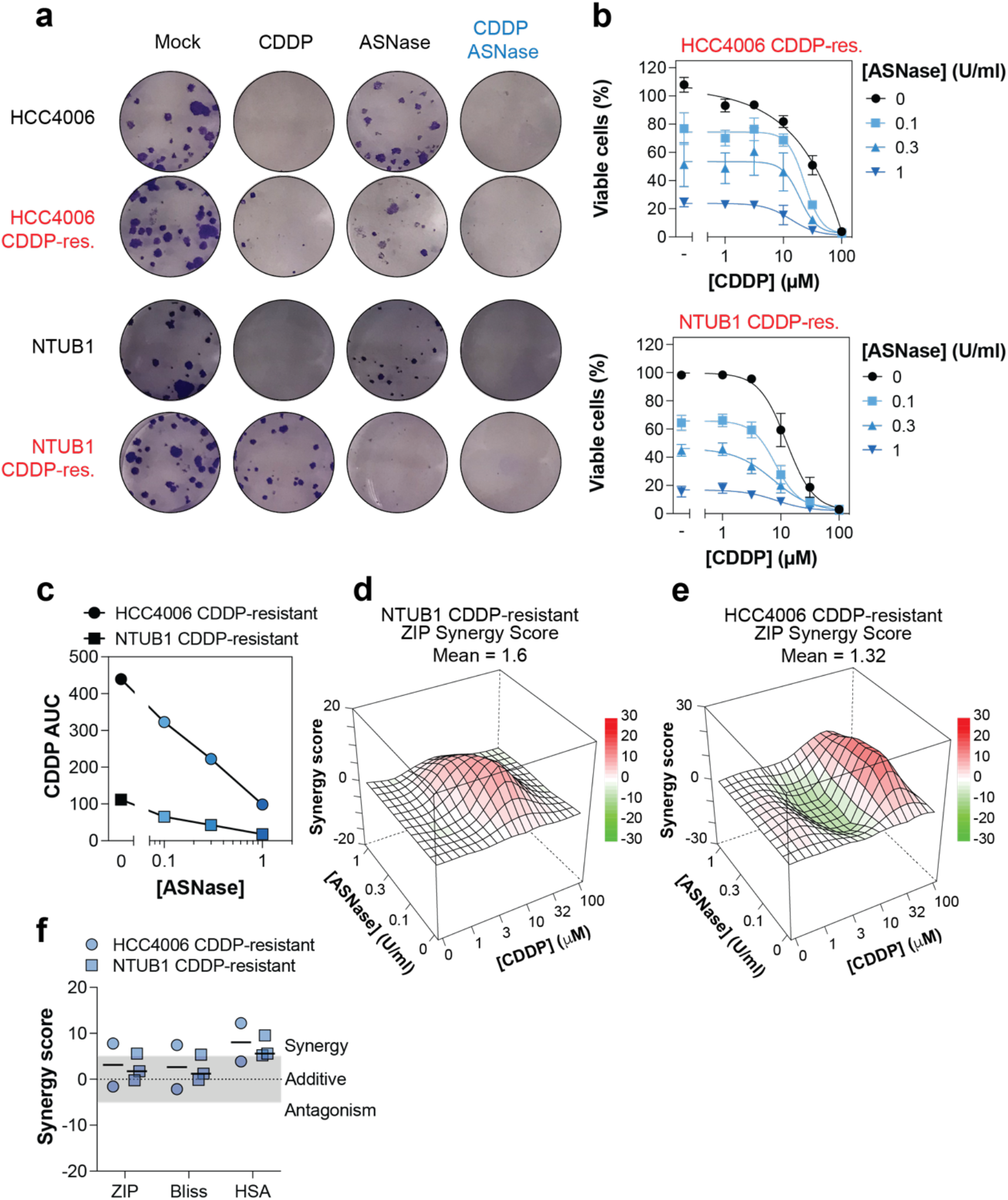
ASNase and cisplatin combination treatment in SLC7A11^high^ cisplatin-resistant cancer cells results in additive inhibition of growth. **a)** Parental and cisplatin (CDDP)-resistant SLC7A11^high^ cell line pairs were cultured in complete media or media supplemented with the indicated therapeutics (0.1 U/ml ASNase, 10 µM CDDP) in a colony outgrowth assay before fixation and staining. Representative images shown (n=2). **b)** CDDP-resistant cell lines were exposed to a CDDP ASNase dose matrix and cell viability measured after 3 days via resazurin reduction. Mean values of independent experiments (n=2-3) each performed in duplicate shown, error bars indicate SEM. **c)** CDDP area under the curve (AUC) as a function of ASNase concentration, derived from the data in (b). Mean values shown (n=2-3). **d, e**) Average synergy landscapes for the indicated cell lines shown, derived from data in (b). **f)** Drug synergy plots for CDDP and ASNase combination in the indicated cell line. Each data point indicates as average delta score from a single dose-response matrix experiment performed in duplicate (n=2-3).

## Discussion

Despite many advances in treatment modalities, cisplatin-containing chemotherapy regimens remain amongst the most widely deployed in the treatment of solid malignancies. One of the main clinical limitations of this therapeutic agent is resistance. Thus, efforts are needed to gain a molecular understanding of the basis of resistance to facilitate the identification of potential biomarkers and treatment strategies to overcome it. In the present study, we used a collection of cancer cell line pairs with acquired resistance to cisplatin, each derived from a different solid malignancy in which cisplatin-based therapy is used (lung, bladder, and ovarian cancer). Using these models, we confirmed a previously reported phenotype of cisplatin-resistant cancer cells, dependence upon extracellular glutamine, and coupled this with upregulation of SLC7A11. Subsequently, we evaluated therapeutic strategies to exploit this acquired nutrient dependency and identified ASNase therapy as a potential actionable strategy.

Metabolic plasticity is a hallmark of cancer that is readily exploited in the acquirement of drug resistance, but this can also lead to collateral vulnerabilities of these drug resistance cells. In the case of acquired resistance to platinum therapy, rewiring of glutamine metabolism can lead to a dependence upon extracellular glutamine, which has been previously documented in cell models [10–16]. Furthermore, cisplatin-resistant tumours grafted in mice have also been shown to be selectively sensitive to nutrient deprivation through repeated fasting cycles compared to cisplatin-sensitive tumours [12], and high GLS1 expression has been correlated with reduced survival of ovarian cancer patients in several independent cohorts [10]. Here, we show that two of the cisplatin-resistant experimental models used in this study, the lung HCC4006 and bladder NTUB1 cell models, also acquire dependence upon extracellular glutamine to sustain cell proliferation. In both these cases, this glutamine addiction could be mechanistically linked to up-regulation of the glutamate-cystine antiporter SLC7A11, as SLC7A11 protein levels were higher in these models and depletion via siRNA could reduce the dependence upon extracellular glutamine. This mechanism is distinct from previous reports documenting glutamine-addiction in cisplatin-resistant cell models, which thus far have been attributed to elevated expression of ASCT2 and/or GLS1 [10,14,16], and promoter methylation-mediated downregulation of GS [11]. Elevated SLC7A11 has been documented in cisplatin resistant cell lines [13,25–27], patient-derived material ex vivo [28], and tumours in vivo [26,27], and in experiments in pre-clinical models, has been demonstrated to contribute to cisplatin resistance via RNAi or targeting with small molecules [13,25–27].

Furthermore, in the case of bladder cancer [26] and gastric cancer [27], high SLC7A11 expression correlates with poorer survival in cisplatin treated patients. We hypothesise that the elevated expression of SLC7A11 we observe in our study seeks to fuel GSH biosynthesis, to facilitate cisplatin resistance, which is consistent with earlier studies [13,25–27]. But a consequence of increased cystine import to fuel GSH biosynthesis is increased glutamate export, which in turn renders these cell models dependent upon glutamine. This is also supported by literature outside the context of chemoresistance that has shown that SLC7A11 can dictate glutamine dependence in cell models [29–32], and is supported in our study by the hypersensitivity of SLC7A11^high^ cells to GLS1 inhibitor CB-839, which indicates the elevated demand for glutamine is to generate glutamate. Taken together, these studies and our own support that SLC7A11 is a chemoresistance factor, functioning via elevation of intracellular drug de-toxifier GSH, and here, we mechanistically link this biomarker of chemoresistance to an acquired metabolic vulnerability, the dependence upon extracellular glutamine.

Linking disease phenotypes with molecular targets is key to the rational design of treatment interventions. As we established the acquired glutamine addiction in cisplatin resistant cells can be a consequence of SLC7A11 up-regulation, we thus next sought to target these SLC7A11^high^ cells. Up-regulation of a target can have differential effects upon cell killing through pharmacological inhibition of that same target. For instance, up-regulation can indicate increased dependence of cells and thus enhanced sensitivity, or conversely, up-regulation can also result in therapy resistance. We first evaluated experimental molecules known to induce ferroptosis that directly target SLC7A11 or downstream processes. We observed that SLC7A11^high^ cisplatin-resistant cells can became cross-resistant to ferroptosis inducers erastin and RSL-3, consistent with overexpression studies outside the context of chemoresistance [33,34], altogether indicating exploiting ferroptosis, and specifically SLC7A11 in SLC7A11^high^ cells, as means to kill cisplatin resistant cells may not be a viable strategy. Next, we evaluated the glutaminase allosteric inhibitor CB-839 (telaglenastat) and observed striking sensitivity in specifically the SLC7A11^high^ cisplatin-resistant models, consistent with an earlier report demonstrating SLC7A11 activity can drive CB-839 sensitivity [30], highlighting a promising context to further investigate the use of this molecule.

ASNase is a clinically approved therapy which was the first to exploit a metabolic dependency in cancer cells, specifically the dependence of acute lymphoblastic leukemic (ALL) cells upon circulating asparagine. However, this enzyme also possesses glutaminase activity and can thus deplete both asparagine and glutamine in plasma [23,24]. Outside the context of chemoresistance, there are several examples of exploiting the glutaminase activity of ASNase for targeting glutamine-dependent cancer cells, including both solid [29,35] and haematological cancer-derived models [36–38]. Although glutamine is highly abundant in plasma [9], ASNase therapy can substantially deplete the level of this amino acid [24], and the glutaminase activity of ASNase is understood to contribute to the anti-leukemic activity but is also considered to be a source of toxicity [24], and deciphering the relative contributions is an active area of research [39]. Given the heterogeneity of glutamine-addiction mechanisms reported in cisplatin-resistant models, whether for nucleotide biosynthesis [12], the TCA cycle [10,16], or the SLC7A11-dependent mechanism reported here, it is clear that whilst the intracellular actionable vulnerabilities may differ, all these chemoresistant models are dependent upon the extracellular abundance of glutamine. For this reason, to overcome the heterogeneity of glutamine-addiction mechanisms, targeting extracellular glutamine via enzymatic depletion by repurposing ASNase therapy or development of a specific glutaminase therapy [40,41], could be a promising approach.

In conclusion, in the present study, we identified an acquired metabolic vulnerability of cisplatin-resistant cancer cells, specifically addiction to glutamine, and mechanistically linked this vulnerability to upregulation of the chemoresistance factor SLC7A11. Subsequently we provide evidence that repurposing of the decades old anti-leukemic therapy ASNase could be a promising approach to target this glutamine addiction, and future studies should interrogate this in more complex pre-clinical models of chemoresistance.

## Materials & methods

### Cell lines

Human lung adenocarcinoma cell line HCC4006 were authenticated (Microsynth) and thereafter cultured in escalating doses of clinical-grade cisplatin (#146262, Hospira Nordic AB) to generate a cell model with acquired resistance. The starting dose of 1 μM cisplatin was raised to 2.5 μM and thereafter 5 μM over a time course of 3 months. HCC4006 cisplatin-resistant cells were subsequently cultured in 5 μM cisplatin for 3 further months. Human bladder cancer cell line NTUB1 and cisplatin resistant derivative NTUB1/P [42] were a gift from Dr Kumar Sanjiv (Karolinska Institutet). Human ovarian cancer A2780 and cisplatin resistant derivative A2780cis were from the European Collection of Authenticated Cell Cultures (ECACC, Sigma-Aldrich). All cell lines were cultured in RPMI 1640 GlutaMAX supplemented with 10% heat-inactivated foetal bovine serum (FBS) and penicillin-streptomycin (100 U/ml and 100 g/ml, respectively). Culture medium was purchased from ThermoFisher Scientific. Glutamine withdrawal experiments were conducted with glutamine-free RPMI 1640 medium (#21870076, ThermoFisher Scientific). Cells were grown at 37 °C in 5% CO_2_ humidified incubators and cultures regularly monitored and tested negative for the presence of mycoplasma using a commercial biochemical test (MycoAlert, Lonza).

### Drugs

Cisplatin (CDDP; Sigma-Aldrich, #P4394) was prepared in 2 mM stock in water or PBS and stored at 4°C. Asparaginase (ASNase, Sigma-Aldrich, #A3809) was prepared in water to a stock concentration of 500 U/ml and stored at −20°C. Erastin (Sigma-Aldrich, #E7781), RSL-3 (SigmaAldrich, #SML2234) and Telaglenastat (CB-839, MedChemExpress, #HY-12248) were prepared in DMSO at a concentration of 10 mM and stored at -20°C.

### Transfections

Transfections were performed using INTERFERin (Polyplus Transfection). SLC7A11 targeting siRNA (Hs_SLC7A11_2 FlexiTube siRNA, #SI00104902, Qiagen) and control siRNA (All Stars Negative Control, Qiagen) were transfected at 10 nM final concentration.

### Immunoblots

Cells were scraped in lysis buffer (50 mM Tris-HCl pH 8, 150 mM NaCl, 1 mM EDTA, 1% Triton X-100, 0.1% SDS) supplemented with cOmplete EDTA-free protease inhibitor cocktail (Roche) and Halt phosphatase inhibitor cocktail (Thermo Fisher). Samples were incubated on ice for 30 mins to 1 hour with occasional vortexing before centrifugation to pellet insoluble material. The Pierce BCA Protein Assay Kit (Thermo Fisher) was used to determine the concentration of the remaining soluble fraction and samples with equal total protein quantity were prepared with Laemmli Sample Buffer (Bio-Rad) before denaturation at 95 °C for 5 min. SDS–PAGE was performed using a Bio-Rad setup with Criterion TGX 4–20% gels (Bio-Rad) and proteins were transferred to a nitrocellulose membrane using Trans-Blot Turbo Transfer System (Bio-Rad), all according to the manufacturer’s instructions. Membranes were blocked in Odyssey Blocking Buffer (Li-Cor) and probed with primary and then species-appropriate IRDye-conjugated secondary antibodies (Li-Cor) before visualisation on an Odyssey Fc Imaging System (Li-Cor). Primary antibodies used in this study: xCT/SLC7A11 (Cell Signaling, #12691S, 1:1000), GLS1 (abcam, #ab156876, 1:2000), GLS2 (abcam, #ab113509, 1:1000), GS (abcam, #ab176562, 1:1000), ASCT2 (Sigma-Aldrich, #ABN73, 1:1000), GPX4 (abcam, #ab125066,1:2000), α-Tubulin (abcam, #ab7291: 1:5000), and GAPDH (Santa Cruz, #32233, 1:1000).

### Proliferation inhibition assays

Inhibition of cell proliferation by monotherapy or combination treatment was determined by resazurin reduction assay as previously described [43]. Compounds prepared either in DMSO (erastin, RSL-3, CB-839; 10 mM) or 0.3% Tween-20 (CDDP, 2mM; ASNase, 250 U/ml) were dispensed in dilution series into clear bottomed 384-well plates (#3764, Corning) using the D300e digital dispenser (Tecan). Solvents were normalised across each plate. Cell suspensions were prepared (50 000 cells/ml) and 50 μl/well dispensed using a MultiDrop (Thermo Fisher) to give a final seeding concentration of 2500 cells/well. Plates were incubated at 37 °C and 5% CO_2_ for 72 h in a humidity chamber until resazurin (#R17017, Sigma Aldrich) dissolved in PBS was added to a final concentration of 0.01 mg/ml. Fluorescence at 530/590 nm (ex/em) was measured after 4-6 h resazurin incubation with Hidex Sense Microplate Reader. Fluorescence intensity for each well was normalised to the average of control wells on the same plate containing cells with solvent (100% viability control) and medium with solvent (0% viability control). The data were analysed using a four-parameter logistic model in Prism 10 (GraphPad Software). For determination of drug combination effects, relative cell viability measurements across a drug matrix were analysed using SynergyFinder+ with ZIP, Bliss, or HSA [44,45]. Synergy summary scores were derived from the average of the synergy scores across the entire dose–response landscape.

### Colony outgrowth assay

Cells (200-800 per well) were seeded into 12-well or 6-well plates. To assess glutamine dependence, cells were seeded in glutamine-free media (#21870076, ThermoFisher Scientific). To assess drug sensitivity, cells were seeded in complete media and 24 h later, media replaced with drug-supplemented media. Following ∼14 days, media were removed, and the wells washed carefully in PBS before the colonies were stained with 4% methylene blue (#M9140, Sigma-Aldrich, Sweden) in methanol for 20 min.

### Statistical analyses

Statistical analyses were conducted using GraphPad Prism 10 software. Specific statistical test details are indicated in the corresponding figure legends. Statistical significance is defined as p < 0.05, unless otherwise stated. Asterisks in figures signifies statistical significance (* for p ≤ 0.05, ** for p ≤ 0.01).

## Author contributions

Conceptualisation, J.W. and S.G.R.; Methodology, J.W., R.S., J.B., S.G.R.; Investigation, J.W. and S.G.R.; Writing – Original Draft, J.W. and S.G.R.; Writing – Reviewing and Editing; J.W. and S.G.R.; Visualisation, J.W. and S.G.R.; Resources, R.S. and J.B.; Funding Acquisition, J.B. and S.G.R.; Supervision, J.B. and S.G.R.

## Availability of data and materials

Data and materials are available from the corresponding author upon request

## Acknowledgements

We thank Sophia Cedar and Klas Wiman (Karolinska Institutet) for sharing reagents for pilot studies and advice.

